# Expression patterns of blood-based biomarkers of neurodegeneration and inflammation across adulthood in rhesus macaques

**DOI:** 10.1101/2024.12.20.629742

**Authors:** Ludwig A.P. Metzler, Jeanette M. Metzger, Keenan J. Gerred, Marina E. Emborg, Amita Kapoor

## Abstract

As the global human population rapidly ages and diseases of aging become more prevalent, preclinical models of age-related neurodegenerative disorders are increasingly important for identifying early diagnostic biomarkers, monitoring disease progression, and evaluating treatment responsiveness. Rhesus macaques are an ideal species for studies on neurodegeneration due to their phylogenetic relatedness to humans and their complex brain anatomy and physiology. Technological advances in assay sensitivity have facilitated the identification of blood-based biomarkers of neurodegeneration and inflammation in human populations. The aim of this study was to translate these methods for use in male and female rhesus macaques across adulthood. We collected plasma samples from 47 rhesus macaques representing pre-adult (1-5 years, n=6 female, n=5 male), young (5-7 years, n=5 female, n=7 male), middle (8-16 years, n=7 female, n=7 male), and older adult (17-22 years, n=6 female, n=4 male) subjects. Quantified biomarkers included neurofilament light chain (NfL), glial fibrillary acidic protein (GFAP), amyloid beta (Aβ42, Aβ40, and their ratio), total tau, phosphorylated tau (pTau181), interleukin (IL) 2, IL-6, IL-8, and IL-10. Plasma NfL and IL-6 levels were significantly increased with age in both sexes, with a marked rise during middle adulthood. The ratio of Aβ42/Aβ40 significantly declined steadily with age, mirroring the aging pattern described in humans. There was no effect of age or sex on total tau or pTau181 levels. Overall, these results demonstrate the feasibility of evaluating blood biomarkers of neurodegeneration and inflammation in rhesus macaques during adulthood.

## 1. INTRODUCTION

The global population of adults aged 65 and older is expected to increase by over 100% in the next 50 years (United Nations Department of Economic and Social Affairs, 2023). With this rise comes a concomitant increase in the prevalence of chronic conditions associated with aging. Research aimed at understanding the aging process is critical, especially for conditions related to neurodegeneration, where early detection can be the key to successful intervention for disease prevention and/or management. Sex differences during aging also play an important role, as women are more affected by chronic conditions, including autoimmune disorders and Alzheimer’s disease (AD) (DuMont et al., 2023). Moreover, women have a longer life expectancy than men, further increasing the lifetime prevalence of disease in females compared to males (Patwardhan et al., 2024).

Blood-based biomarkers of neurodegeneration have long been an objective of aging research to measure key pathophysiological processes for screening, diagnosis, progression, and treatment monitoring. They are envisioned as a replacement of biomarkers in cerebral spinal fluid (CSF), offering a less invasive approach and vastly improving their clinical utility (Dark et al., 2024). These assays are now a reality due to major technological advances in sensitivity and reproducibility, which allow measurement of these biomarkers at levels at least 10 times lower than found in CSF with a high degree of correlation (Ashton et al., 2020; Leuzy et al., 2022).

The key target blood-based, age-related biomarkers that have been studied in humans include neurofilament light chain (NfL), glial fibrillary acidic protein (GFAP), the (amyloid beta) Aβ42/40 ratio, tau proteins, and inflammatory cytokines. NfL supports neuronal cytoskeletal structure and, if axonal damage occurs, is released into extracellular space and can then move into CSF and blood. As such, NfL is currently considered a non-specific marker of neuronal damage and its levels in blood and CSF have been shown to increase in humans with age and with Alzheimer’s disease (AD) progression (Gaetani et al., 2019; Simonsen et al., 2023). GFAP is the main intermediate filament protein in the cytoskeleton of mature astrocytes (Middeldorp and Hol, 2011). Although the mechanism by which GFAP is released from astrocytes is not fully elucidated, it can be measured in the CSF and blood and has been approved for use as a blood biomarker for diagnosis of mild traumatic brain injury (Abdelhak et al., 2022). Similar to NfL, GFAP is a non-specific biomarker of neurodegeneration and its levels have been reported to increase with age in blood of humans and in the brains of humans and rats (Nichols et al., 1993; Tybirk et al., 2023).

Aβ and tau deposition are core pathological components of AD and can be observed in other forms of dementia (Mehta and Schneider, 2023). Aβ is a byproduct of the amyloid precursor protein (APP) proteolytic cleavage, APP processing can generate the more soluble form Aβ1-40 and the insoluble Aβ1-42, which is more likely to accumulate in the walls of the cerebrocortical and leptomeningeal blood vessels (Zecca et al., 2021).The Aβ42/40 ratio reflects a balance between the two peptides, and provides an indirect measure of amyloid plaque deposition and AD progression. A decreased ratio is linked to a higher risk of developing AD in humans; additionally, the Aβ42/40 ratio has been shown to decrease with age in both human and rhesus macaques (Pérez-Grijalba et al., 2019; Zhao et al., 2017). The protein tau stabilizes microtubules in neurons; however, when tau becomes hyperphosphorylated, it can accumulate, aggregate, and form neurofibrillary tangles that affect neuronal homeostasis (Thijssen et al., 2021). The tau protein can undergo various post translation modifications, including phosphorylation at multiple sites, resulting in the presence of differing forms of phosphorylated tau (pTau) (Thijssen et al., 2021). Elevated pTau181 has been shown to be associated with cognitive decline in humans (Coomans et al., 2023). Additionally, total tau levels, which combine the measurement of all forms of the tau protein, have also been associated with AD and other neurodegenerative conditions (Pillai et al., 2019).

Cytokines are also promising blood-based indicators of age-related changes. Indeed, it has long been established that the inflammatory milieu increases with age (Li et al., 2023). Pro-inflammatory cytokines play a variety of roles and are produced by many tissue types. For example, they are produced in response to tissue injury and aid in immune cell recruitment. IL-6 and IL-8 are elevated in neurodegenerative conditions and are associated with worsened cognitive function (Clark and Peterson, 1994; Maggio et al., 2006). In contrast, anti-inflammatory cytokines such as IL-10 help counteract the effects of inflammation and play a protective role. In human studies, anti-inflammatory cytokines can prevent age-related inflammation (Dagdeviren et al., 2017).

Rhesus macaques (*Macaca mulatta*) are commonly used as models for human aging and age-related diseases due to their phylogenetic relatedness, similar physiology, and complex neuroanatomy (Beckman et al., 2021). Further, they have similar sex-based patterns, with females living longer than males (Clochero et al., 2016). Particularly relevant for the current study, rhesus macaques display age-related deposition of Aβ in the brain parenchyma and vasculature (Heuer et al., 2012; Uno, 1993). Moreover, pTau fibrils have been detected in the entorhinal cortex and hippocampus of aged rhesus macaques (Datta et al., 2021; Härtig et al., 2000; Paspalas et al., 2018). The blood-based biomarkers typically evaluated in aging humans have not been studied in the normally aging rhesus macaque. In rhesus CSF, Robertson and colleagues reported an age-related decline in Aβ40 and Aβ42, but not in NfL, total tau, or pTau181; males had higher levels of NfL and total tau compared to females (Robertson et al., 2022). The aim of the present study was to evaluate expression patterns of blood-based biomarkers of neurodegeneration and inflammation across adulthood in male and female rhesus macaques as these physiological changes are proposed to start years, if not decades, before disease diagnosis (Greaves and Rohrer, 2019). This will provide insights into the biological aging processes and assess the validity of the rhesus as a preclinical model for human aging research.

## 2. METHODS

### 2.1. Animals

All procedures were performed in accordance with the NIH guide for the Care and Use of Laboratory Animals and under the approval of the Vice Chancellor Office for Research and Graduate Education Institutional Animal Care and Use Committee. Animals were housed in enclosures that meet the requirements specified in the Animal Welfare Act Regulations and the Guide for the Care and Use of Laboratory Animals Guide. All animals were evaluated by veterinary staff or trained animal care staff at least twice daily for signs of pain, distress, and illness.

The rhesus macaques (*Macaca mulatta*) used in this study were cross-sectional samples of convenience from the colony at the Wisconsin National Primate Research Center (WNPRC). A total of 47 rhesus macaques were selected: 24 female and 23 male. Groups were further subdivided by age with animals designated as pre-adults (1-5 years old), young adults (5-7 years), middle-aged (8-16 years) and older adults (17-22 years). The division of age groups in this study was based on previously published studies (Kulik et al., 2015; Simmons, 2016). The population demographics are presented in Table 1.

**Table 1.** Demographic information for study. All animals used in the study were rhesus macaques (*Macaca mulatta*). Posthoc statistical significance is shown as follows; ^a^ p < 0.05 compared to other marked age groups. ^b^ p < 0.05 compared to other marked age groups. c p < 0.05 compared to all groups.

| Experimental Group | Pre-adult | Young adult | Middle-aged adults | Old adult |
| --- | --- | --- | --- | --- |
| Age Range of Group | 1-5 years | 5-7 years | 8-16 years | 17-22 years |
| Age; years; mean ± SD<br>(Min-Max)<br>All | 3.66 ± 0.75<br>(1.66 - 4.96) | 6.16 ± 0.84<br>(5.00 - 7.63) | 11.98 ± 2.88<br>(8.10 - 16.97) | 19.01 ± 1.27<br>(17.39 - 22.39) |
| Female | 3.80 ± 0.26<br>(3.33 - 4.07) | 5.97 ± 0.72<br>(5.31 - 7.22) | 12.36 ± 2.89<br>(8.10 - 16.97) | 18.85 ± 0.52<br>(18.30 - 19.62) |
| Male | 3.50 ± 1.06<br>(1.66 - 4.96) | 6.30 ± 0.89<br>(5.00 - 7.63) | 11.59 ± 2.83<br>(8.52 - 16.25) | 19.25 ± 1.88<br>(17.39 - 22.39) |
| Number |  |  |  |  |
| Female | 6 | 5 | 7 | 6 |
| Male | 5 | 7 | 7 | 4 |
| Weight in kg |  |  |  |  |
| Female (mean ± SD) | 5.16 ± 0.42<sup>a</sup> | 6.24 ± 1.44<sup>a</sup> | 10.21 ± 2.13<sup>a</sup> | 8.19 ± 1.76 |
| Male (mean ± SD) | 5.42 ± 1.89<sup>b</sup> | 9.24 ± 1.33<sup>b</sup> | 14.86 ± 3.71<sup>b</sup> | 12.51 ± 0.99<sup>b</sup> |
| Female parity (n) | N/A | 0 | 1 | 7<sup>c</sup> |

Rhesus macaques were sedated for blood draws with ketamine and their vital signs monitored until recovered from anesthesia. Blood was collected in vacutainer tubes for EDTA plasma, kept on ice and centrifuged within 15 minutes of collection due to instability some of the analytes of interest (Sunde et al. 2023). All plasma samples were stored at -80°C until assay.

### 2.2. Assays

Mesoscale Discovery (MSD, Maryland, USA) electrochemiluminescent (ECL) multiplexed immunoassay kits were used for the measurement of Aβ40, Aβ42, IFN-γ, IL-1β, IL-2, IL-6, IL-8, IL-10, GFAP, NF-L, total Tau, and pT181 in rhesus macaque EDTA plasma. The kits are sandwich assays which utilize arrays of up to 10 different capture antibodies coated on spots on a conductive circuit in the wells of a 96 well plate, and detection antibodies conjugated to an ECL tag. An MSD QuickPlex SQ 120MM ECL plate imager controlled by a computer running MSD Methodical Mind software was used to read the plates, and assay data was calculated in MSD Discovery Workbench 4.0 using 4PL curve fits. The catalog and analyte information for the kits used are presented in Table S1, and all were performed as per the kit protocols without modification. In the Amyloid Beta Peptide Panel, Aβ40 Blocker was omitted as the levels of Aβ40 in EDTA plasma samples are known to be relatively low, and the blocker is only required when running samples with high expected Aβ40 levels. Analytes that were below the limit of detection for most samples were excluded from analysis (Aβ38, IFN-γ, IL-1β).

### 2.3. Statistical analysis

Data was assessed for normality using a Shapiro-Wilk test (Shapiro and Wilk, 1965), and outliers were identified through Tukey’s fences (Schwertman et al., 2004). For statistical analysis, outliers were winsorized by replacing them with the 95th and 5th percentile values for high and low, respectively (Schoonjans et al., 2011). IL-2 was not detectable in 6 out of 36 samples, therefore the lower limit of detection of the assay was substituted. Summary statistics of the raw dataset (without winsorization) are presented in Tables S2 and S3. To determine the effect of age-group and sex on individual biomarker levels and weight, we used a two-way ANOVA with interaction (age-group, sex, age-group x sex). If a significant interaction was found, analysis was further conducted using 1-way ANOVA. The effect of age on parity in females was assessed by 1-way ANOVA. Multiple comparisons were conducted with Tukey’s Test. A p-value of <0.05 was considered significant for all analyses performed during this study. Normality and outlier assessments were performed through R (v4.4.1; R Core Team, v3.5.1; ggplot2), while all other statistical tests were performed using GraphPad Prism (Version 10.4.1).

## 3. RESULTS

### 3.1. Demographics

Animal weight changed non-linearly with age (F(3, 39) = 24.26, p < 0.0001) and was affected by sex (F(1, 39) = 23, p < 0.0001). In both sexes, weight steadily increased from pre-adult to middle-age but decreased in both sexes at the oldest age group. As expected, parity significantly increased with age as none of the young adult females in this cohort had given birth (F(2, 15) = 16, p = 0.0002) (Table 1).

### 3.2. Neurodegeneration biomarkers

NfL levels increased significantly with age in both sexes (Figure 1A; F(3,39) = 4.374, p = 0.0095), with the oldest animals showing markedly higher levels compared to the pre-adult (p = 0.0146) and young adult groups (p = 0.0317).

**Figure 1.**
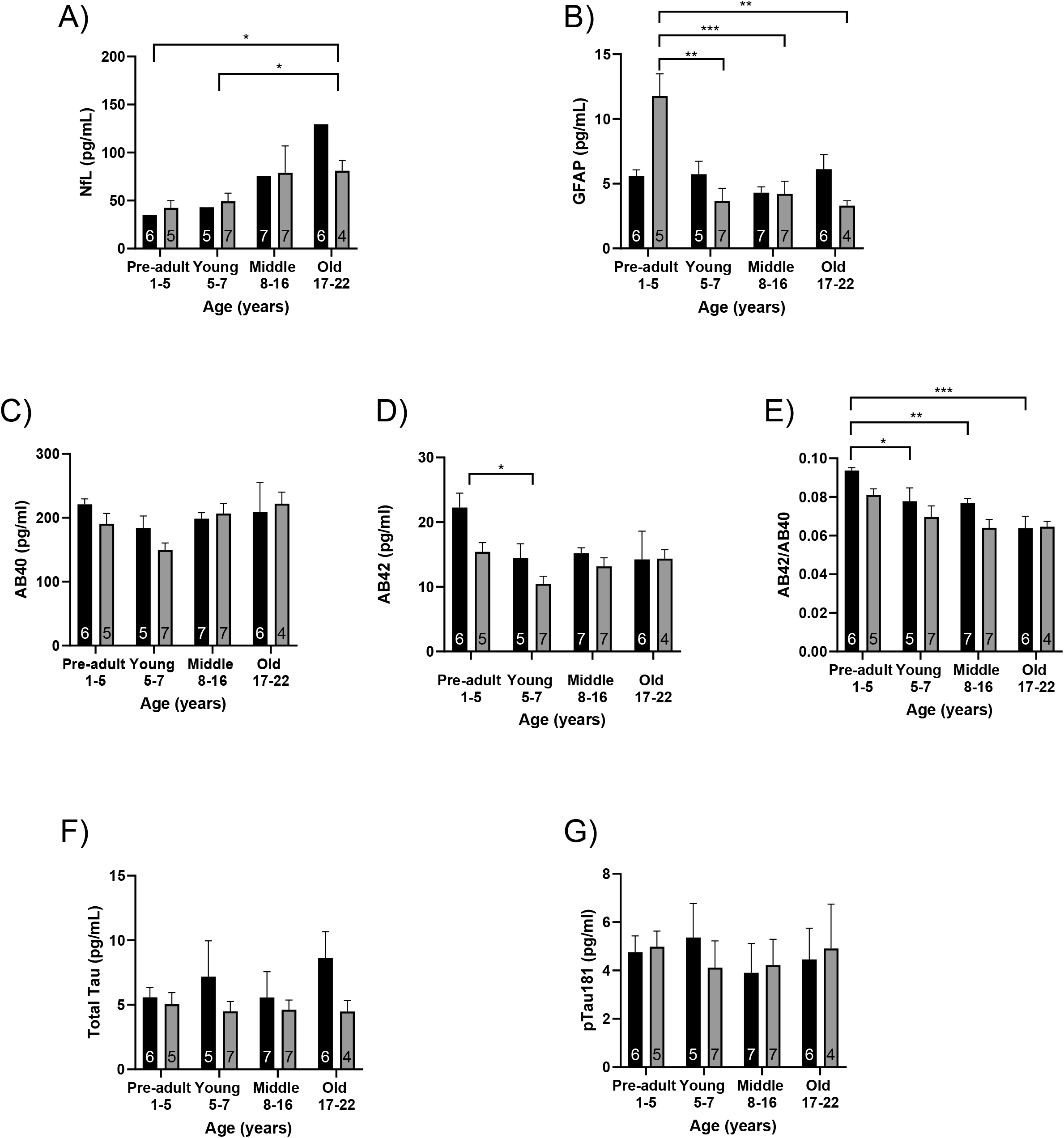
Blood expression levels of biomarkers (mean ± SEM) across age and sex in females (black bars) and males (grey bars); *n* is indicated within each bar. The biomarkers shown are (A) NfL, (B) GFAP, (C) Aβ42, (D) Aβ40, (E) Aβ42/Aβ40 ratio, (F) pTau181, and (G) total tau. Posthoc statistical significance is shown; * p<0.05, ** p<0.01, *** p<0.001 for age comparisons.

Plasma GFAP levels showed a significant age x sex interaction (Figure 1B; F(3, 39) = 8.112, p = 0.0003). Further, sex-stratified analysis showed a significant effect of age in males only (F = 11.24, p = 0.0002) with higher GFAP levels in pre-adults compared to young adults (p = 0.0004), middle-aged adults (p = 0.0008), and older adults (p = 0.0010). Females did not have any age-related changes in plasma GFAP levels.

Age and sex did not influence plasma A⍰40 (Figure 1C). However, both age and sex affected A⍰42 levels (Figure 1D). There was a significant decrease in plasma A⍰42 levels with age (F(3, 39) = 3.117 , p = 0.0370) and higher levels overall in females (F(1, 39) = 4.342 , p = 0.0438) which was driven by high levels of plasma A⍰42 in the pre-adult females. The A⍰42 to A⍰40 ratio (Figure 1E) showed the expected aging pattern for A⍰ isoform changes, with a significant decline with age (F(3, 39) = 8.015, p = 0.0003), and females had higher levels of this ratio compared to males (F(1, 39) = 5.965, p = 0.0192). Further posthoc analysis showed that the pre-adult females had a higher ratio compared to middle-aged (p=0.0484) and older females (p=0.0003).

Plasma levels of total tau (Figure 1F) or pTau181 (Figure 1G) were not affected by age or sex. All neurodegeneration biomarker statistics are presented in Table S4.

#### 3.3. Cytokines

A trend towards an effect of age was detected on plasma levels of IL-2 (Figure 2A; F(3, 39) = 2.267, p = 0.0959) with lower levels of IL-2 between the young adult and oldest adult groups (p=0.0602). In contrast, there was a clear effect of increasing plasma IL-6 concentrations with age (Figure 2B; F(3, 39) = 7.577, p = 0.0004). IL-6 levels dramatically increased in middle-aged adults compared to pre-adult (p=0.0037) and young adult (p=0.0016) animals. This was also reflected in the young adult to older adult comparison (p=0.0466). There were no significant effects of age, sex, or age-group interaction on IL-8 (Figure 2C) or IL-10 levels (Figure 2D). All statistics for inflammatory biomarkers are presented in Table S5.

**Figure 2.**
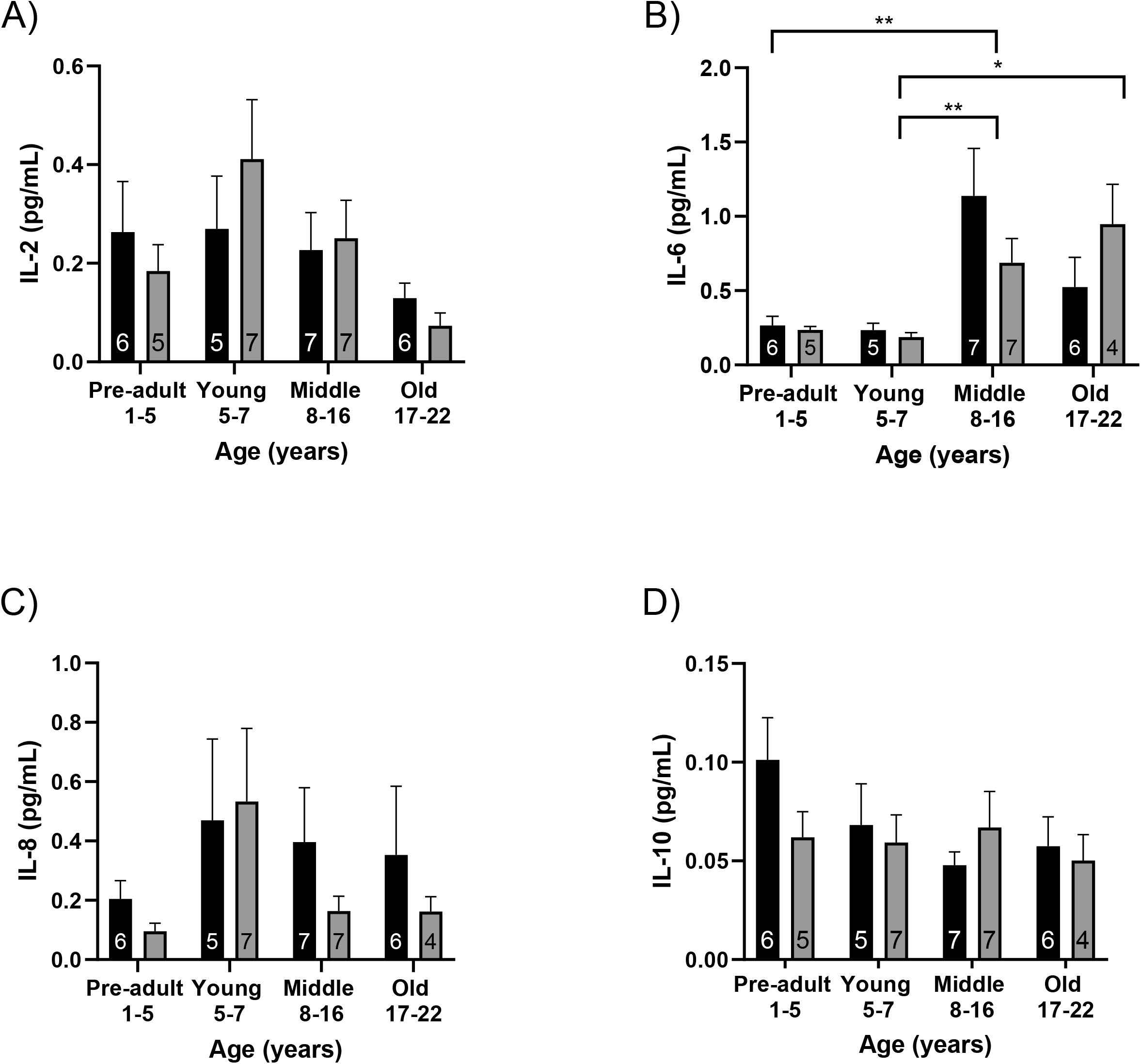
Blood expression levels of cytokines (mean ± SEM) across age and sex in females (black bars) and males (grey bars); *n* is indicated within each bar. The cytokines shown are (A) IL-2, (B) IL-6, (C) IL-8, and (D) IL-10. Posthoc statistical significance is shown; * p<0.05, ** p<0.01, *** p<0.001 for age comparisons.

## 4. DISCUSSION

The results of this study demonstrate the feasibility of evaluating blood biomarkers of neurodegeneration and inflammation in rhesus macaques across adulthood. Ultra-sensitive methodology to conduct these measurements has only recently become available; this report is the first to show successful measure of these analytes using the commercially available assays from MSD with ECL technology in rhesus macaque blood.

NfL blood levels were significantly increased during middle adulthood (8-17 years), which coincided with an increase in IL-6 levels, suggesting a shift in the rhesus macaque physiology at this age. To the best of our knowledge, only one other report has evaluated NfL levels although in the CSF of rhesus macaque and found no association with age (Robertson et al., 2022). It should be noted that in the aforementioned study, the oldest animal was 12 years old, which is the mean of the middle-aged group in the current study. An increase in circulating NfL levels with age was also reported in common marmoset monkeys (Phillips et al., 2024). In normally aging humans, age-dependent increases in blood levels of NfL have been documented (Arslan and Zetterberg, 2023). Interestingly, higher NfL levels in cognitively normal humans were associated with worse cognitive performance at baseline and accelerated cognitive decline in a follow-up visit. In these same participants, higher NfL levels were also associated with MRI measures of diminished white matter integrity, volume of white matter hyperintensities, and presence of lacunes (van Arendonk et al., 2024).

GFAP levels were higher in male pre-adult compared to adult rhesus, likely as a result of higher astrocytic activity due to rapid brain development (Tybirk et al., 2023). Indeed, higher levels of GFAP during childhood have been demonstrated in human populations in both sexes (Tybirk et al., 2023). Although we aimed to match age and sex, the youngest macaques in our study were males (1.66 years for males vs. 3.33 years for females). In adults, GFAP levels have consistently been higher in female patients with AD (Abbas et al., 2023), traumatic brain injury (Sass et al., 2021), and Parkinson’s disease (Li et al., 2023). This sex difference has also been replicated in other mammals such as rats, where GFAP-immunoreactivity was higher in the female brain (Rasia-Filho et al., 2002). A higher astrocyte reactivity in females compared to males has been proposed to be a factor in the higher incidence of AD in women (Abbas et al., 2023), however, we did not observe this pattern in our study. This discrepancy may be attributed to the unexpectedly high levels of plasma GFAP observed in pre-adult males, which likely obscured statistical differences in the adult group during our two-way ANOVA analyses.

We did not identify an effect of age or sex on plasma A⍰40 levels, yet there were significant decreases with age in A⍰42 levels and A⍰42/A⍰40 ratio. The decrease in A⍰42 levels was unexpected and was driven by the high levels of A⍰42 in pre-adult females. In humans, the decrease in the A⍰42/A⍰40 ratio is due to a change in the predominant isoform of A⍰ shifting to A⍰42. Indeed, low plasma levels of the A⍰42/40 ratio were associated with more pronounced decline in cognitive function (Giudici et al., 2020), and similar findings have been replicated in a number of studies (Graff-Radford et al., 2007; Okereke et al., 2009). One other report in rhesus macaques have shown no change in age-related CSF levels of A⍰40 or A⍰42, and, similar to the current study, the concentrations of A⍰40 were higher than A⍰42, as it is the predominant isoform (Appleman et al., 2024).

In normally aging humans, levels of total plasma tau were reported to be higher in elderly compared to middle-aged adults (Chiu et al., 2017) and were associated with decreased performance in memory tests (Cantero et al., 2020). Specifically, pTau181 plasma levels were negatively correlated with gray matter volume (Tissot et al., 2021). Lastly, high levels of pTau181 have been demonstrated in AD and are proposed for differential diagnoses of AD with frontemporal dementia (Meng and Lei, 2020; Thijssen et al., 2021). The relationship between pathological pTau accumulation in the brain of rhesus macaques and resultant circulating levels have not yet been explored. Follow up studies are warranted, including quantitative analyses of other forms of phosphorylated tau, such as pTau217, due to its close relationship to measures of amyloid and tau deposition by brain imaging with positron emission tomography (Thijssen et al., 2021).

There is a wealth of evidence from human population studies showing that with increasing age, there is chronic low-grade inflammation, a weak response to novel antigens, and poor immune recall responses (Wong et al., 2021). Immunosenescence has been cited as the root cause of the increased incidence and severity of infectious diseases, cancers, and autoimmune diseases observed to occur with age (Mittelbrunn and Kroemer, 2021; Pawelec, 2020). IL-6 is a cytokine that is produced by cells of the immune system, as well as vascular endothelial cells, adipocytes, and skeletal muscle. IL-6 is sometimes referred to as the ‘geriatric biomarker’ due to its robust association with disease, disability, and mortality (Singh and Newman, 2011). The data from the current study supports this aging phenotype in the rhesus macaque, as there was a significant increase in IL-6 levels between the young and middle-aged adults which maintained through adulthood. Our data narrows the age window to 8-17 years, as the period when the chronic inflammation begins, and, as mentioned above, this is also the age where NfL levels rise. Similar to the results in the current study, one other report in found that there was an age-related increase of markers of inflammation in rhesus macaques aged 15 to >20 years (Walker et al., 2019).

IL-2 is a growth factor that promotes natural killer cell activity and the differentiation of naïve T cells into Th1 and Th2 cells. It can also act to negatively regulate IL-17 production, which is linked to neutrophil mobilization and T cell recruitment (Rea et al., 2009; Zenobia and Hajishengallis, 2015). Studies in humans have demonstrated that IL-2 is decreased in the 8^th^ and 9^th^ decade of life (Rea et al., 2009; Whisler et al., 1996). Our results are in accordance with those reports, as IL-2 trended towards being lower in the old-adult group compared to the young rhesus. These data could also mean that the T cells of the rhesus macaque exhibit diminished generation of IL-2 earlier in the lifespan compared to humans. There is one paper that is contradictory to our data that showed plasma IL-2 increased with age in rhesus macaques; however, the study, which was performed over a decade earlier utilized xMAP, which is less sensitive analytical platform (Didier et al., 2012). The precise mechanisms underlying the increase in inflammatory markers during adulthood are unknown. However, some pro-inflammatory, age-related changes could be increased total and visceral adiposity, declining levels of sex hormones, and oxidative damage (Klein and Flanagan, 2016; Martínez de Toda et al., 2023).

As part of the animals’ demographics, we collected data on the animals’ weights. We identified a significant decline in weight in older adults compared to middle-aged rhesus that mirrored the age of muscle decline previously seen in dual-energy X-ray absorptiometry (DXA) analysis in rhesus (Colman et al. 2005). Sarcopenia, the universal, involuntary decline in skeletal muscle mass and function with age, is also observed in humans and has been reported to account for approximately 50% decline from peak levels by the 8^th^ decade of life (Larsson et al., 2019).

The majority of plasma levels of the biomarkers in this study were similar to those obtained in human studies that utilized the MSD platform, except for GFAP and A⍰42, which were approximately 10 times lower in rhesus macaques (Kivisäkk et al., 2023; Spanos et al., 2022; Ulndreaj et al., 2023). Others have found that different species have varying levels of similar blood biomarkers. For example, Robertson et al., detected higher in rhesus macaque CSF levels of A⍰40, A⍰42, and NfL compared to cynomolgus macaques (Robertson et al., 2022). To our knowledge, there are no publications that have measured plasma levels of A⍰s in humans with the MSD platform, but concentrations in the current study were similar to those obtained by liquid chromatography-tandem mass spectrometry (Weber et al., 2024). There is currently a lack of standardized reference material published for these analytes. This paucity of data should be considered when comparing raw data across different studies for absolute concentrations and age reference ranges. However, levels of NfL were significantly correlated with data generated via Simoa, an alternative analytical method for measurement of low levels of biomarkers (Ulndreaj et al., 2023). As interest in measurement of these biomarkers is rapidly increasing, we expect that more work toward standardization and/or conversion factors will continue (Arslan and Zetterberg, 2023).

The major limitation of this report was the lack of macaques considered elderly. Average longevity of rhesus macaques is 25 years, but the oldest known macaque lived to age 41 in captivity (Alldritt et al., 2024). In our study, animals of ages 17-22 years were considered older adults, relative to the other monkeys the study population. We used a cross-sectional study design that utilized samples of convenience from the colony at the WNPRC. Future studies will benefit from employing longitudinal design to determine the rate of change of these biomarkers in individuals, as well as including rhesus macaques over 22 years of age.

Overall, this study has generated foundational data on blood biomarkers of neurodegeneration and inflammation across rhesus macaques’ adulthood. We have identified age-related changes in rhesus blood samples that mirror findings in the human population. These findings provide a starting point for future studies on the progression and prevention of neurodegenerative changes, offering a valuable comparative perspective to bridge preclinical and clinical research.

## Supporting information

These are supplemental Tables, referenced in the Manuscript

## 5. ACKNOWLEDGEMENTS

We gratefully acknowledge the technical assistance of the WNPRC Assay Services Unit and Animal Care staff. We also thank Eric Peterson and Kerri Fuchs from the WNPRC Scientific Protocol Implementation unit for their assistance in collection of samples. This research was supported by NIH awards R61/R33 NS115102, R01NS124857, P51OD011106, UL1TR002373, and the Rainwater Foundation.

## Notes

### Competing Interest Statement

The authors have declared no competing interest.

## REFERENCES

Abbas, S., Bellaver, B., Ferreira, P.C.L., Povala, G., Ferrari-Souza, J.P., Lussier, F.Z., Zalzale, H., Lemaire, P.C., Soares, C., Rohden, F., Aguzzoli, C.S., Cabrera, A., Leffa, D.T., Therriault, J., Tissot, C., Servaes, S., Macedo, A.C., Chamoun, M., Bezgin, G., Stevenson, J., Stevenson, A., Rahmouni, N., Pallen, V., Karikari, T.K., Ashton, N.J., Benedet, A.L., Zetterberg, H., Blennow, K., Rosa-Neto, P., Pascoal, T.A., 2023. Sex impacts the association of plasma Glial Fibrillary Acidic Protein with neurodegeneration in Alzheimer’s disease. Alzheimers Dement. 19, e076402. 10.1002/alz.076402

Abdelhak, A., Foschi, M., Abu-Rumeileh, S., Yue, J.K., D’Anna, L., Huss, A., Oeckl, P., Ludolph, A.C., Kuhle, J., Petzold, A., Manley, G.T., Green, A.J., Otto, M., Tumani, H., 2022. Blood GFAP as an emerging biomarker in brain and spinal cord disorders. Nat. Rev. Neurol. 18, 158–172. 10.1038/s41582-021-00616-3

Alldritt, S., Ramirez, J.S.B., Wael, R.V. de, Bethlehem, R., Seidlitz, J., Wang, Z., Nenning, K., Esper, N.B., Smallwood, J., Franco, A.R., Byeon, K., Alexander-Bloch, A., Amaral, D.G., Amiez, C., Balezeau, F., Baxter, M.G., Becker, G., Bennett, J., Berkner, O., Blezer, E.L.A., Brambrink, A.M., Brochier, T., Butler, B., Campos, L.J., Canet-Soulas, E., Chalet, L., Chen, A., Cléry, J., Constantinidis, C., Cook, D.J., Dehaene, S., Dorfschmidt, L., Drzewiecki, C.M., Erdman, J.W., Everling, S., Falchier, A., Fleysher, L., Fox, A., Freiwald, W., Froesel, M., Froudist-Walsh, S., Fudge, J., Funck, T., Gacoin, M., Gale, D.J., Gallivan, J., Garin, C.M., Griffiths, T.D., Guedj, C., Hadj-Bouziane, F., Hamed, S.B., Harel, N., Hartig, R., Hiba, B., Howell, B.R., Jarraya, B., Jung, B., Kalin, N., Karpf, J., Kastner, S., Klink, C., Kovacs-Balint, Z.A., Kroenke, C., Kuchan, M.J., Kwok, S.C., Lala, K.N., Leopold, D.A., Li, G., Lindenfors, P., Linn, G., Mars, R.B., Masiello, K., Menon, R.S., Messinger, A., Meunier, M., Mok, K., Morrison, J.H., Nacef, J., Nagy, J., Neudecker, V., Neuringer, M., Noonan, M.P., Ortiz-Rios, M., Perez-Zoghbi, J.F., Petkov, C.I., Pinsk, M., Poirier, C., Procyk, E., Rajimehr, R., Reader, S.M., Rudko, D.A., Rushworth, M.F.S., Russ, B.E., Sallet, J., Sanchez, M.M., Schmid, M.C., Schwiedrzik, C.M., Scott, J.A., Sein, J., Sharma, K.K., 2024. Brain Charts for the Rhesus Macaque Lifespan. bioRxiv 2024.08.28.610193. 10.1101/2024.08.28.610193

Appleman, M.-L., Thomas, J.L., Weiss, A.R., Nilaver, B.I., Cervera-Juanes, R., Kohama, S.G., Urbanski, H.F., 2024. Effect of hormone replacement therapy on amyloid beta (Aβ) plaque density in the rhesus macaque amygdala. Front. Aging Neurosci. 15, 1326747. 10.3389/fnagi.2023.1326747

Arslan, B., Zetterberg, H., 2023. Neurofilament light chain as neuronal injury marker – what is needed to facilitate implementation in clinical laboratory practice? Clin. Chem. Lab. Med. CCLM 61, 1140–1149. 10.1515/cclm-2023-0036

Ashton, N.J., Hye, A., Rajkumar, A.P., Leuzy, A., Snowden, S., Suárez-Calvet, M., Karikari, T.K., Schöll, M., La Joie, R., Rabinovici, G.D., Höglund, K., Ballard, C., Hortobágyi, T., Svenningsson, P., Blennow, K., Zetterberg, H., Aarsland, D., 2020. An update on blood-based biomarkers for non-Alzheimer neurodegenerative disorders. Nat. Rev. Neurol. 16, 265–284. 10.1038/s41582-020-0348-0

Beckman, D., Chakrabarty, P., Ott, S., Dao, A., Zhou, E., Janssen, W.G., Donis-Cox, K., Muller, S., Kordower, J.H., Morrison, J.H., 2021. A novel tau-based rhesus monkey model of Alzheimer’s pathogenesis. Alzheimers Dement. J. Alzheimers Assoc. 17, 933–945. 10.1002/alz.12318

Cantero, J.L., Atienza, M., Ramos-Cejudo, J., Fossati, S., Wisniewski, T., Osorio, R.S., 2020. Plasma tau predicts cerebral vulnerability in aging. Aging 12, 21004–21022. 10.18632/aging.104057

Chiu, M.-J., Fan, L.-Y., Chen, T.-F., Chen, Y.-F., Chieh, J.-J., Horng, H.-E., 2017. Plasma Tau Levels in Cognitively Normal Middle-Aged and Older Adults. Front. Aging Neurosci. 9. 10.3389/fnagi.2017.00051

Clark, J.A., Peterson, T.C., 1994. Cytokine production and aging: overproduction of IL-8 in elderly males in response to lipopolysaccharide. Mech. Ageing Dev. 77, 127–139. 10.1016/0047-6374(94)90020-5

Clochero, F., Rau, R., Jones, O.R., Barthold, J.A., Conde, D.A., 2016. The emergence of longevous populations [WWW Document]. 10.1073/pnas.1612191113

Coomans, E.M., Verberk, I.M.W., Ossenkoppele, R., Verfaillie, S.C.J., Visser, D., Gouda, M., Tuncel, H., Wolters, E.E., Timmers, T., Windhorst, A.D., Golla, S.S.V., Scheltens, P., van, W.M., der Flier, van Berckel, B.N.M., Teunissen, C.E., 2023. A Head-to-Head Comparison Between Plasma pTau181 and Tau PET Along the Alzheimer’s Disease Continuum. J. Nucl. Med. 64, 437–443. 10.2967/jnumed.122.264279

Dagdeviren, S., Jung, D.Y., Friedline, R.H., Noh, H.L., Kim, J.H., Patel, P.R., Tsitsilianos, N., Inashima, K., Tran, D.A., Hu, X., Loubato, M.M., Craige, S.M., Kwon, J.Y., Lee, K.W., Kim, J.K., 2017. IL-10 prevents aging-associated inflammation and insulin resistance in skeletal muscle. FASEB J. 31, 701–710. 10.1096/fj.201600832R

Dark, H.E., Duggan, M.R., Walker, K.A., 2024. Plasma biomarkers for Alzheimer’s and related dementias: A review and outlook for clinical neuropsychology. Arch. Clin. Neuropsychol. Off. J. Natl. Acad. Neuropsychol. 39, 313–324. 10.1093/arclin/acae019

Datta, D., Leslie, S.N., Wang, M., Morozov, Y.M., Yang, S., Mentone, S., Zeiss, C., Duque, A., Rakic, P., Horvath, T.L., van Dyck, C.H., Nairn, A.C., Arnsten, A.F.T., 2021. Age-related calcium dysregulation linked with tau pathology and impaired cognition in non-human primates. Alzheimers Dement. J. Alzheimers Assoc. 17, 920–932. 10.1002/alz.12325

Didier, E.S., Sugimoto, C., Bowers, L.C., Khan, I.A., Kuroda, M.J., 2012. Immune correlates of aging in outdoor-housed captive rhesus macaques (Macaca mulatta). Immun. Ageing 9, 25. 10.1186/1742-4933-9-25

DuMont, M., Agostinis, A., Singh, K., Swan, E., Buttle, Y., Tropea, D., 2023. Sex representation in neurodegenerative and psychiatric disorders’ preclinical and clinical studies. Neurobiol. Dis. 184, 106214. 10.1016/j.nbd.2023.106214

Gaetani, L., Blennow, K., Calabresi, P., Filippo, M.D., Parnetti, L., Zetterberg, H., 2019. Neurofilament light chain as a biomarker in neurological disorders. J. Neurol. Neurosurg. Psychiatry 90, 870– 881. 10.1136/jnnp-2018-320106

Giudici, K.V., de Souto Barreto, P., Guyonnet, S., Li, Y., Bateman, R.J., Vellas, B., MAPT/DSA Group, 2020. Assessment of Plasma Amyloid-β42/40 and Cognitive Decline Among Community-Dwelling Older Adults. JAMA Netw. Open 3, e2028634. 10.1001/jamanetworkopen.2020.28634

Graff-Radford, N.R., Crook, J.E., Lucas, J., Boeve, B.F., Knopman, D.S., Ivnik, R.J., Smith, G.E., Younkin, L.H., Petersen, R.C., Younkin, S.G., 2007. Association of Low Plasma Aβ42/Aβ40 Ratios With Increased Imminent Risk for Mild Cognitive Impairment and Alzheimer Disease. Arch. Neurol. 64, 354–362. 10.1001/archneur.64.3.354

Greaves, C.V., Rohrer, J.D., 2019. An update on genetic frontotemporal dementia. J. Neurol. 266, 2075–2086. 10.1007/s00415-019-09363-4

Härtig, W., Klein, C., Brauer, K., Schüppel, K.F., Arendt, T., Brückner, G., Bigl, V., 2000. Abnormally phosphorylated protein tau in the cortex of aged individuals of various mammalian orders. Acta Neuropathol. (Berl.) 100, 305–312. 10.1007/s004010000183

Heuer, E., Rosen, R.F., Cintron, A., Walker, L.C., 2012. Nonhuman primate models of Alzheimer-like cerebral proteopathy. Curr. Pharm. Des. 18, 1159–1169. 10.2174/138161212799315885

Kivisäkk, P., Carlyle, B.C., Sweeney, T., Trombetta, B.A., LaCasse, K., El-Mufti, L., Tuncali, I., Chibnik, L.B., Das, S., Scherzer, C.R., Johnson, K.A., Dickerson, B.C., Gomez-Isla, T., Blacker, D., Oakley, D.H., Frosch, M.P., Hyman, B.T., Aghvanyan, A., Bathala, P., Campbell, C., Sigal, G., Stengelin, M., Arnold, S.E., 2023. Plasma biomarkers for diagnosis of Alzheimer’s disease and prediction of cognitive decline in individuals with mild cognitive impairment. Front. Neurol. 14. 10.3389/fneur.2023.1069411

Klein, S.L., Flanagan, K.L., 2016. Sex differences in immune responses. Nat. Rev. Immunol. 16, 626– 638. 10.1038/nri.2016.90

Kulik, L., Amici, F., Langos, D., Widdig, A., 2015. Sex Differences in the Development of Social Relationships in Rhesus Macaques (Macaca mulatta). Int. J. Primatol. 36, 353–376. 10.1007/s10764-015-9826-4

Larsson, L., Degens, H., Li, M., Salviati, L., Lee, Y.I., Thompson, W., Kirkland, J.L., Sandri, M., 2019. Sarcopenia: Aging-Related Loss of Muscle Mass and Function. Physiol. Rev. 99, 427–511. 10.1152/physrev.00061.2017

Leuzy, A., Mattsson-Carlgren, N., Palmqvist, S., Janelidze, S., Dage, J.L., Hansson, O., 2022. Blood-based biomarkers for Alzheimer’s disease. EMBO Mol. Med. 14, e14408. 10.15252/emmm.202114408

Li, X., Li, C., Zhang, W., Wang, Y., Qian, P., Huang, H., 2023. Inflammation and aging: signaling pathways and intervention therapies. Signal Transduct. Target. Ther. 8, 1–29. 10.1038/s41392-023-01502-8

Maggio, M., Guralnik, J.M., Longo, D.L., Ferrucci, L., 2006. Interleukin-6 in Aging and Chronic Disease: A Magnificent Pathway. J. Gerontol. A. Biol. Sci. Med. Sci. 61, 575–584.

Martínez de Toda, I., González-Sánchez, M., Díaz-Del Cerro, E., Valera, G., Carracedo, J., Guerra-Pérez, N., 2023. Sex differences in markers of oxidation and inflammation. Implications for ageing. Mech. Ageing Dev. 211, 111797. 10.1016/j.mad.2023.111797

Mehta, R.I., Schneider, J.A., 2023. Neuropathology of the Common Forms of Dementia. Clin. Geriatr. Med. 39, 91–107. 10.1016/j.cger.2022.07.005

Meng, J., Lei, P., 2020. Plasma pTau181 as a biomarker for Alzheimer’s disease. MedComm 1, 74–76. 10.1002/mco2.1

Middeldorp, J., Hol, E.M., 2011. GFAP in health and disease. Prog. Neurobiol. 93, 421–443. 10.1016/j.pneurobio.2011.01.005

Mittelbrunn, M., Kroemer, G., 2021. Hallmarks of T cell aging. Nat. Immunol. 22, 687–698. 10.1038/s41590-021-00927-z

Nichols, N.R., Day, J.R., Laping, N.J., Johnson, S.A., Finch, C.E., 1993. GFAP mRNA increases with age in rat and human brain. Neurobiol. Aging 14, 421–429. 10.1016/0197-4580(93)90100-p

Okereke, O.I., Xia, W., Selkoe, D.J., Grodstein, F., 2009. Ten-Year Change in Plasma Amyloid β Levels and Late-Life Cognitive Decline. Arch. Neurol. 66, 1247–1253. 10.1001/archneurol.2009.207

Paspalas, C.D., Carlyle, B.C., Leslie, S., Preuss, T.M., Crimins, J.L., Huttner, A.J., van Dyck, C.H., Rosene, D.L., Nairn, A.C., Arnsten, A.F.T., 2018. The aged rhesus macaque manifests Braak stage III/IV Alzheimer’s-like pathology. Alzheimers Dement. J. Alzheimers Assoc. 14, 680–691. 10.1016/j.jalz.2017.11.005

Patwardhan, V., Gil, G.F., Arrieta, A., Cagney, J., DeGraw, E., Herbert, M.E., Khalil, M., Mullany, E.C., O’Connell, E.M., Spencer, C.N., Stein, C., Valikhanova, A., Gakidou, E., Flor, L.S., 2024. Differences across the lifespan between females and males in the top 20 causes of disease burden globally: a systematic analysis of the Global Burden of Disease Study 2021. Lancet Public Health 9, e282–e294. 10.1016/S2468-2667(24)00053-7

Pawelec, G., 2020. The human immunosenescence phenotype: does it exist? Semin. Immunopathol. 42, 537–544. 10.1007/s00281-020-00810-3

Pérez-Grijalba, V., Romero, J., Pesini, P., Sarasa, L., Monleón, I., San-José, I., Arbizu, J., Martínez-Lage, P., Munuera, J., Ruiz, A., Tárraga, L., Boada, M., Sarasa, M., 2019. Plasma Aβ42/40 Ratio Detects Early Stages of Alzheimer’s Disease and Correlates with CSF and Neuroimaging Biomarkers in the AB255 Study. J. Prev. Alzheimers Dis. 6, 34–41. 10.14283/jpad.2018.41

Phillips, K.A., Lopez, M., Bartling-John, E., Meredith, R., Buteau, A., Alvarez, A., Ross, C.N., 2024. Serum biomarkers associated with aging and neurodegeneration in common marmosets (Callithrix jacchus). Neurosci. Lett. 819, 137569. 10.1016/j.neulet.2023.137569

Pillai, J.A., Bonner-Jackson, A., Bekris, L.M., Safar, J., Bena, J., Leverenz, J.B., 2019. Highly Elevated Cerebrospinal Fluid Total Tau Level Reflects Higher Likelihood of Non-Amnestic Subtype of Alzheimer’s Disease. J. Alzheimers Dis. JAD 70, 1051–1058. 10.3233/JAD-190519

Rasia-Filho, A.A., Xavier, L.L., dos Santos, P., Gehlen, G., Achaval, M., 2002. Glial fibrillary acidic protein immunodetection and immunoreactivity in the anterior and posterior medial amygdala of male and female rats. Brain Res. Bull. 58, 67–75. 10.1016/S0361-9230(02)00758-X

Rea, I.M., Stewart, M., Campbell, P., Alexander, H.D., Crockard, A.D., Morris, T.C.M., 2009. Changes in Lymphocyte Subsets, Interleukin 2, and Soluble Interleukin 2 Receptor in Old and Very Old Age. Gerontology 42, 69–78. 10.1159/000213775

Robertson, E.L., Boehnke, S.E., Lyra E Silva, N. de M., Armitage-Brown, B., Winterborn, A., Cook, D.J., De Felice, F.G., Munoz, D.P., 2022. Characterization of cerebrospinal fluid biomarkers associated with neurodegenerative diseases in healthy cynomolgus and rhesus macaque monkeys. Alzheimers Dement. N. Y. N 8, e12289. 10.1002/trc2.12289

Sass, D., Guedes, V.A., Smith, E.G., Vorn, R., Devoto, C., Edwards, K.A., Mithani, S., Hentig, J., Lai, C., Wagner, C., Dunbar, K., Hyde, D.R., Saligan, L., Roy, M.J., Gill, J., 2021. Sex Differences in Behavioral Symptoms and the Levels of Circulating GFAP, Tau, and NfL in Patients With Traumatic Brain Injury. Front. Pharmacol. 12, 746491. 10.3389/fphar.2021.746491

Schoonjans, F., De Bacquer, D., Schmid, P., 2011. Estimation of population percentiles. Epidemiol. Camb. Mass 22, 750–751. 10.1097/EDE.0b013e318225c1de

Schwertman, N.C., Owens, M.A., Adnan, R., 2004. A simple more general boxplot method for identifying outliers. Comput. Stat. Data Anal. 47, 165–174. 10.1016/j.csda.2003.10.012

Shapiro, S.S., Wilk, M.B., 1965. An analysis of variance test for normality (complete samples)†. Biometrika 52, 591–611. 10.1093/biomet/52.3-4.591

Simmons, H.A., 2016. Age-Associated Pathology in Rhesus Macaques (Macaca mulatta). Vet. Pathol. 53, 399–416. 10.1177/0300985815620628

Simonsen, A.H., Gleerup, H.S., Musaeus, C.S., Sellebjerg, F., Hansen, M.B., Søndergaard, H.B., Waldemar, G., Hasselbalch, S.G., 2023. Neurofilament light chain levels in serum among a large mixed memory clinic cohort: Confounders and diagnostic usefulness. Alzheimers Dement. Diagn. Assess. Dis. Monit. 15, e12512. 10.1002/dad2.12512

Singh, T., Newman, A.B., 2011. Inflammatory markers in population studies of aging. Ageing Res. Rev. 10, 319–329. 10.1016/j.arr.2010.11.002

Spanos, M., Shachar, S., Sweeney, T., Lehmann, H.I., Gokulnath, P., Li, G., Sigal, G.B., Nagaraj, R., Bathala, P., Rana, F., Shah, R.V., Routenberg, D.A., Das, S., 2022. Elevation of neural injury markers in patients with neurologic sequelae after hospitalization for SARS-CoV-2 infection. iScience 25, 104833. 10.1016/j.isci.2022.104833

Thijssen, E.H., Joie, R.L., Strom, A., Fonseca, C., Iaccarino, L., Wolf, A., Spina, S., Allen, P.I.E., Cobigo, Y., Heuer, H., VandeVrede, L., Proctor, N.K., Lago, A.L., Baker, S., Sivasankaran, R., Kieloch, A., Kinhikar, A., Yu, L., Valentin, M.-A., Jeromin, A., Zetterberg, P.H., Hansson, P.O., Mattsson-Carlgren, N., Graham, D., Blennow, P.K., Kramer, P.J.H., Grinberg, L.T., Seeley, P.W.W., Rosen, P.H., Boeve, P.B.F., Miller, P.B.L., Teunissen, P.C.E., Rabinovici, P.G.D., Rojas, J.C., Dage, J.L., Boxer, P.A.L., 2021. Association of Plasma P-tau217 and P-tau181 with clinical phenotype, neuropathology, and imaging markers in Alzheimer’s disease and frontotemporal lobar degeneration: a retrospective diagnostic performance study. Lancet Neurol. 20, 739. 10.1016/S1474-4422(21)00214-3

Tissot, C. L., Benedet, A., Therriault, J., Pascoal, T.A., Lussier, F.Z., Saha-Chaudhuri, P., Chamoun, M., Savard, M., Mathotaarachchi, S.S., Bezgin, G., Wang, Y.-T., Fernandez Arias, J., Rodriguez, J.L., Snellman, A., Ashton, N.J., Karikari, T.K., Blennow, K., Zetterberg, H., De Villers-Sidani, E., Huot, P., Gauthier, S., Rosa-Neto, P., for the Alzheimer’s Disease Neuroimaging Initiative, 2021. Plasma pTau181 predicts cortical brain atrophy in aging and Alzheimer’s disease. Alzheimers Res. Ther. 13, 69. 10.1186/s13195-021-00802-x

Tybirk, L., Hviid, C.V.B., Knudsen, C.S., Parkner, T., 2023. Serum GFAP – pediatric reference interval in a cohort of Danish children. Clin. Chem. Lab. Med. CCLM 61, 2041–2045. 10.1515/cclm-2023-0280

Ulndreaj, A., Sohaei, D., Thebault, S., Pons-Belda, O.D., Fernandez-Uriarte, A., Campbell, C., Cheo, D., Stengelin, M., Sigal, G., Freedman, M.S., Scarisbrick, I.A., Prassas, I., Diamandis, E.P., n.d. Quantitation of neurofilament light chain protein in serum and cerebrospinal fluid from patients with multiple sclerosis using the MSD R-PLEX NfL assay. Diagn. Berl. Ger. 10, 275–280. 10.1515/dx-2022-0125

United Nations Department of Economic and Social Affairs, 2023. World Social Report 2023: Leaving No One Behind in an Ageing World, World Social Report. United Nations. 10.18356/9789210019682

Uno, H., 1993. The incidence of senile plaques and multiple infarction in aged macaque brain. Neurobiol. Aging 14, 673–674. 10.1016/0197-4580(93)90067-l

van Arendonk, J., Wolters, F.J., Neitzel, J., Vinke, E.J., Vernooij, M.W., Ghanbari, M., Ikram, M.A., 2024. Plasma neurofilament light chain in relation to 10-year change in cognition and neuroimaging markers: a population-based study. GeroScience 46, 57–70. 10.1007/s11357-023-00876-5

Walker, E.M., Slisarenko, N., Gerrets, G.L., Kissinger, P.J., Didier, E.S., Kuroda, M.J., Veazey, R.S., Jazwinski, S.M., Rout, N., 2019. Inflammaging phenotype in rhesus macaques is associated with a decline in epithelial barrier-protective functions and increased pro-inflammatory function in CD161-expressing cells. GeroScience 41, 739–757. 10.1007/s11357-019-00099-7

Weber, D.M., Taylor, S.W., Lagier, R.J., Kim, J.C., Goldman, S.M., Clarke, N.J., Vaillancourt, D.E., Duara, R., McFarland, K.N., Wang, W., Golde, T.E., Racke, M.K., 2024. Clinical utility of plasma Aβ42/40 ratio by LC-MS/MS in Alzheimer’s disease assessment. Front. Neurol. 15. 10.3389/fneur.2024.1364658

Whisler, R.L., Beiqing, L., Chen, M., 1996. Age-Related Decreases in IL-2 Production by Human T Cells Are Associated with Impaired Activation of Nuclear Transcriptional Factors AP-1 and NF-AT. Cell. Immunol. 169, 185–195. 10.1006/cimm.1996.0109

Wong, G.C.L., Ng, T.K.S., Lee, J.L., Lim, P.Y., Chua, S.K.J., Tan, C., Chua, M., Tan, J., Lee, S., Sia, A., Ng, M.K.W., Mahendran, R., Kua, E.H., Ho, R.C.M., Larbi, A., 2021. Horticultural Therapy Reduces Biomarkers of Immunosenescence and Inflammaging in Community-Dwelling Older Adults: A Feasibility Pilot Randomized Controlled Trial. J. Gerontol. A. Biol. Sci. Med. Sci. 76, 307–317. 10.1093/gerona/glaa271

Zecca, C., Pasculli, G., Tortelli, R., Dell’Abate, M.T., Capozzo, R., Barulli, M.R., Barone, R., Accogli, M., Arima, S., Pollice, A., Brescia, V., Logroscino, G., 2021. The Role of Age on Beta-Amyloid1–42 Plasma Levels in Healthy Subjects. Front. Aging Neurosci. 13, 698571. 10.3389/fnagi.2021.698571

Zenobia, C., Hajishengallis, G., 2015. Basic biology and role of interleukin-17 in immunity and inflammation. Periodontol. 2000 69, 142–159. 10.1111/prd.12083

Zhao, Q., Lu, J., Yao, Z., Wang, S., Zhu, L., Wang, J., Chen, B., 2017. Upregulation of Aβ42 in the Brain and Bodily Fluids of Rhesus Monkeys with Aging. J. Mol. Neurosci. MN 61, 79–87. 10.1007/s12031-016-0840-6

