## Supplementary material for "Expression patterns of blood-based biomarkers of neurodegeneration and inflammation across adulthood in rhesus macaques": These are supplemental Tables, referenced in the Manuscript

Table S1. Mesoscale Discovery (MSD) kits used for biomarker analysis in rhesus macaques. Amyloid beta (Aβ), interferon (IFN), interleukin (IL), glial fibrillary acidic protein (GFAP), neurofilament light chain (NfL), phosphorylated Tau (pTau).

| **MSD Kit** | **Catalog Number** | **Analytes** | **Protocol Version** | **Dilution Factor**  **(µL sample:µL buffer)** |
| --- | --- | --- | --- | --- |
| V-PLEX Plus Amyloid Beta (Aβ) Peptide Panel 1 4G8 Human | K15199E | Aβ38, Aβ40, Aβ42 | 18084-v5-2019Jan | 4X (20:60) |
| V-PLEX Plus Proinflammatory Panel 1 NHP | K15056D | IFN-𝛾, IL-1β, IL-2, IL-6, IL-8, IL-10 | 18115-v4-2018Feb | 2X (60:60) |
| S-PLEX Neurology Panel 1 NHP | K15640S | GFAP, NfL, Total Tau Total | 18275-v2-2023Sep | 2X (40:40) |
| S-PLEX Tau (pT181) NHP | K156AGMS | pTau181 | 18269-v1-2021Dec | None (70:0) |

Table S2. Summary statistics for raw (un-winsorized) analysis of all biomarkers across all ages.

| **Analyte** | **Sex** | **Range (Min - Max) (pg/mL)** | **Mean  (pg/mL)** | **SD  (pg/mL)** |
| --- | --- | --- | --- | --- |
| **NfL** | All | 18.456 - 617.391 | 76.291 | 93.476 |
|  | Female | 18.456 - 617.391 | 90.143 | 122.455 |
|  | Male | 20.238 - 208.564 | 61.837 | 42.416 |
| **GFAP** | All | 1.798 - 163.346 | 8.849 | 23.039 |
|  | Female | 2.012 - 163.364 | 11.800 | 31.645 |
|  | Male | 1.798 - 17.063 | 5.769 | 4.601 |
| **Ratio Aβ42/40** | All | 0.041 - 0.099 | 0.074 | 0.014 |
|  | Female | 0.045 - 0.099 | 0.078 | 0.015 |
|  | Male | 0.041 - 0.092 | 0.070 | 0.012 |
| **Aβ40** | All | 102.263 -435.511 | 196.348 | 52.035 |
|  | Female | 136.128 - 435.511 | 203.884 | 58.415 |
|  | Male | 102.263 - 275.795 | 188.484 | 43.026 |
| **Aβ42** | All | 5.730 - 34.855 | 14.845 | 5.706 |
|  | Female | 6.084 - 34.855 | 16.577 | 3.467 |
|  | Male | 5.730 - 20.626 | 13.038 | 6.787 |
| **Total Tau** | All | 1.922 - 24.163 | 5.981 | 4.460 |
|  | Female | 2.163 - 24.163 | 7.047 | 5.557 |
|  | Male | 1.922 - 13.517 | 4.868 | 2.450 |
| **pTau181** | All | 1.020 - 27.729 | 4.791 | 4.317 |
|  | Female | 1.020 - 27.729 | 5.471 | 5.729 |
|  | Male | 2.074 - 7.966 | 4.080 | 1.687 |

Table S3. Summary statistics for raw (un-winsorized) analysis of all cytokines across all ages.

| **Cytokine** | **Group** | **Range (Min - Max) (pg/mL)** | **Mean (pg/mL)** | **SD (pg/mL)** |
| --- | --- | --- | --- | --- |
| **IL - 2** | All | 0.028 - 0.920 | 0.245 | 0.233 |
|  | Female | 0.028 - 0.920 | 0.240 | 0.225 |
|  | Male | 0.028 - 0.895 | 0.258 | 0.242 |
| **IL - 6** | All | 0.060 - 4.065 | 0.596 | 0.755 |
|  | Female | 0.079 - 2.584 | 0.597 | 0.669 |
|  | Male | 0.060 - 4.065 | 0.595 | 0.835 |
| **IL - 8** | All | 0.030 - 7.059 | 0.409 | 1.039 |
|  | Female | 0.030 - 1.460 | 0.314 | 0.371 |
|  | Male | 0.032 - 7.059 | 0.508 | 1.429 |
| **IL - 10** | All | 0.010 - 0.179 | 0.064 | 0.038 |
|  | Female | 0.013 - 0.132 | 0.065 | 0.041 |
|  | Male | 0.010 - 0.179 | 0.064 | 0.035 |

Supplementary Table 4. ANOVA results for normalized and winsorized neurodegeneration biomarkers

|  | **NfL** | | **GFAP** | | **Aβ ratio** | | **Aβ40** | | **Aβ42** | | **total Tau** | | **pTau181** | |
| --- | --- | --- | --- | --- | --- | --- | --- | --- | --- | --- | --- | --- | --- | --- |
|  | F(DFn, DFd) | p | F(DFn, DFd) | p | F(DFn, DFd) | p | F(DFn, DFd) | p | F(DFn, DFd) | p | F(DFn, DFd) | p | F(DFn, DFd) | p |
| **Age** | F(3, 39) = 4.374 | 0.0095 | F(3, 39) = 8.814 | 0.0001 | F(3, 39) = 8.015 | 0.0003 | F(3, 39) = 1.909 | 0.1442 | F(3, 39) = 3.117 | 0.0370 | F(3, 39) = 0.344 | 0.7938 | F(3, 39) = 0.210 | 0.8890 |
| **Sex** | F(1, 39) = 0.314 | 0.5785 | F(1, 39) = 0.183 | 0.6711 | F(1, 39) = 5.965 | 0.0192 | F(1, 39) = 0.524 | 0.4735 | F(1, 39) = 4.342 | 0.0438 | F(1, 39) = 3.591 | 0.0655 | F(1, 39) = 0.005 | 0.9426 |
| **Interaction** | F(3, 39) = 0.826 | 0.4874 | F(3, 39) = 8.112 | 0.0003 | F(3, 39) = 0.831 | 0.4849 | F(3, 39) = 0.666 | 0.5782 | F(3, 39) = 0.867 | 0.4666 | F(3, 39) = 0.544 | 0.6553 | F(3, 39) = 0.227 | 0.8768 |

Supplementary Table 5) Anova results for normalized and winsorized data for the analysis of Cytokines.

|  | **IL - 6** | | **IL - 6** | | **IL - 8** | | **IL - 10** | |
| --- | --- | --- | --- | --- | --- | --- | --- | --- |
|  | F(DFn, DFd) | p | F(DFn, DFd) | p | F(DFn, DFd) | p | F(DFn, DFd) | p |
| **Age** | F(3, 39) = 2.267 | 0.0959 | F(3, 39) = 7.577 | 0.0004 | F(3, 39) = 1.308 | 0.2855 | F(3, 39) = 1.104 | 0.3590 |
| **Sex** | F(1, 39) = 0.014 | 0.9059 | F(1, 39) = 0.039 | 0.8455 | F(1, 39) = 0.825 | 0.3693 | F(1, 39) = 0.619 | 0.4361 |
| **Interaction** | F(3, 39) = 0.623 | 0.6047 | F(3, 39) = 1.848 | 0.1545 | F(3, 39) = 0.274 | 0.8442 | F(3, 39) = 1.163 | 0.3363 |
